# Multi-Omics factor analysis - a framework for unsupervised integration of multi-omic data sets

**DOI:** 10.1101/217554

**Authors:** Ricard Argelaguet, Britta Velten, Damien Arnol, Sascha Dietrich, Thorsten Zenz, John C. Marioni, Wolfgang Huber, Florian Buettner, Oliver Stegle

**Affiliations:** European Molecular Biology Laboratory, European Bioinformatics Institute, Hinxton, Cambridge, UK; European Molecular Biology Laboratory (EMBL), Heidelberg, Germany.; Cancer Research UK Cambridge Institute, University of Cambridge, Cambridge, UK; Heidelberg University Hospital, Heidelberg, Germany; German Cancer Research Center (dkfz) and National Center for Tumor Diseases (NCT), Heidelberg, Germany & Hematology, University Hospital Zurich and University of Zurich, 8091 Zurich, Switzerland; Wellcome Trust Sanger Institute, Hinxton, Cambridge, UK; Helmholtz Zentrum München–German Research Center for Environmental Health, Institute of Computational Biology, Neuherberg, Germany

## Abstract

Multi-omic studies promise the improved characterization of biological processes across molecular layers. However, methods for the unsupervised integration of the resulting heterogeneous datasets are lacking. We present Multi-Omics Factor Analysis (MOFA), a computational method for discovering the principal sources of variation in multi-omic datasets. MOFA infers a set of (hidden) factors that capture biological and technical sources of variability. It disentangles axes of heterogeneity that are shared across multiple modalities and those specific to individual data modalities. The learnt factors enable a variety of downstream analyses, including identification of sample subgroups, data imputation, and the detection of outlier samples. We applied MOFA to a cohort of 200 patient samples of chronic lymphocytic leukaemia, profiled for somatic mutations, RNA expression, DNA methylation and *ex-vivo* drug responses. MOFA identified major dimensions of disease heterogeneity, including immunoglobulin heavy chain variable region status, trisomy of chromosome 12 and previously underappreciated drivers, such as response to oxidative stress. In a second application, we used MOFA to analyse single-cell multiomics data, identifying coordinated transcriptional and epigenetic changes along cell differentiation.

## Introduction

Technological advances increasingly enable multiple biological layers to be probed in parallel, ranging from genome, epigenome, transcriptome, proteome and metabolome to phenome profiling (Hasin et al, 2017). Integrative analyses that use information across these data modalities promise to deliver more comprehensive insights into the biological systems under study. Motivated by this, multi-omic profiling is increasingly applied across biological domains, including cancer biology (Cancer Genome Atlas Research Network, 2017; Gerstung et al, 2015; Iorio et al, 2016; Mertins et al, 2016), regulatory genomics (Chen et al, 2016), microbiology (Kim et al, 2016) or host-pathogen biology (Soderholm et al, 2016). Most recent technological advances have also enabled performing multi-omics analyses at the single cell level (Angermueller et al, 2016; Clark et al, 2018; Colomé-Tatché & Theis, 2018; Guo et al, 2017; Macaulay et al, 2015). A common aim of such applications is to characterize heterogeneity between samples, as manifested in one or several of the omic data types (Ritchie et al, 2015). Multi-omics profiling is particularly appealing if the relevant axes of variation are not known *a priori*, and hence may be missed by studies that consider a single data modality or targeted approaches.

A basic strategy for the integration of omics data is testing for marginal associations between different data modalities. A prominent example is molecular QTL-analysis, where large numbers of association tests are performed between individual genetic variants and gene expression levels (Consortium, 2015) or epigenetic marks (Chen et al, 2016). While eminently useful for variant annotation, such association studies are inherently *local* and do not provide a coherent global map of the molecular differences between samples. A second strategy is the use kernel- or graph-based methods to combine different data types into a common similarity network between samples (Lanckriet et al, 2004; Wang et al, 2014); however, it is difficult to pinpoint the molecular determinants of the resulting graph structure. Related to this, there exist generalizations of other clustering methods to reconstruct discrete groups of samples based on multiple data modalities (Mo et al, 2013; Shen et al, 2009).

A key challenge that is not sufficiently addressed by these approaches is interpretability. In particular, it would be desirable to reconstruct the underlying factors that drive the observed variation across samples, similar to the loadings in conventional principal component analysis. These could be continuous gradients, discrete clusters, or combinations thereof. Such factors would also help in establishing or explaining associations with external data such as phenotypes or clinical covariates. Although factor models that aim to address this have previously been proposed, e.g., (Meng et al, 2014; Meng et al, 2016; Singh et al, 2016; Tenenhaus et al, 2014), these methods either lack sparsity, which can reduce interpretability, or they require a substantial number of parameters to be determined in extensive cross-validation or post hoc. Further challenges faced by existing methods are computational scalability to larger datasets, handling of missing values and non-Gaussian data modalities, such as binary readouts or count-based traits.

## Results

We present Multi-Omics Factor Analysis (MOFA), a statistical method for integrating multiple modalities of omic data in an unsupervised fashion. Intuitively, MOFA can be viewed as a versatile and statistically rigorous generalization of principal component analysis (PCA) to multi-omics data. Given several data matrices with measurements of multiple ‘omics data types on the same or on partially overlapping sets of samples, MOFA infers an interpretable low-dimensional data representation in terms of (hidden) factors. These learnt factors represent the driving sources of variation across data modalities, thus facilitating the identification of molecular states or subgroups of samples. The inferred factor loadings are sparse, thereby facilitating the linkage between the factors and its driving molecular features. Importantly, MOFA disentangles to what extent each of the factors is unique to a single data modality or is manifested in multiple modalities (**Fig. 1**), thereby identifying links between the different ‘omics layers. Once trained, the model output can be used for a range of downstream analyses, including visualisation, clustering and classification of samples in the low-dimensional space(s) spanned by the factors as well as automated annotation of factors using (gene set) enrichment analysis, the identification of outlier samples and the imputation of missing values (**Fig. 1**).

**Figure 1:**
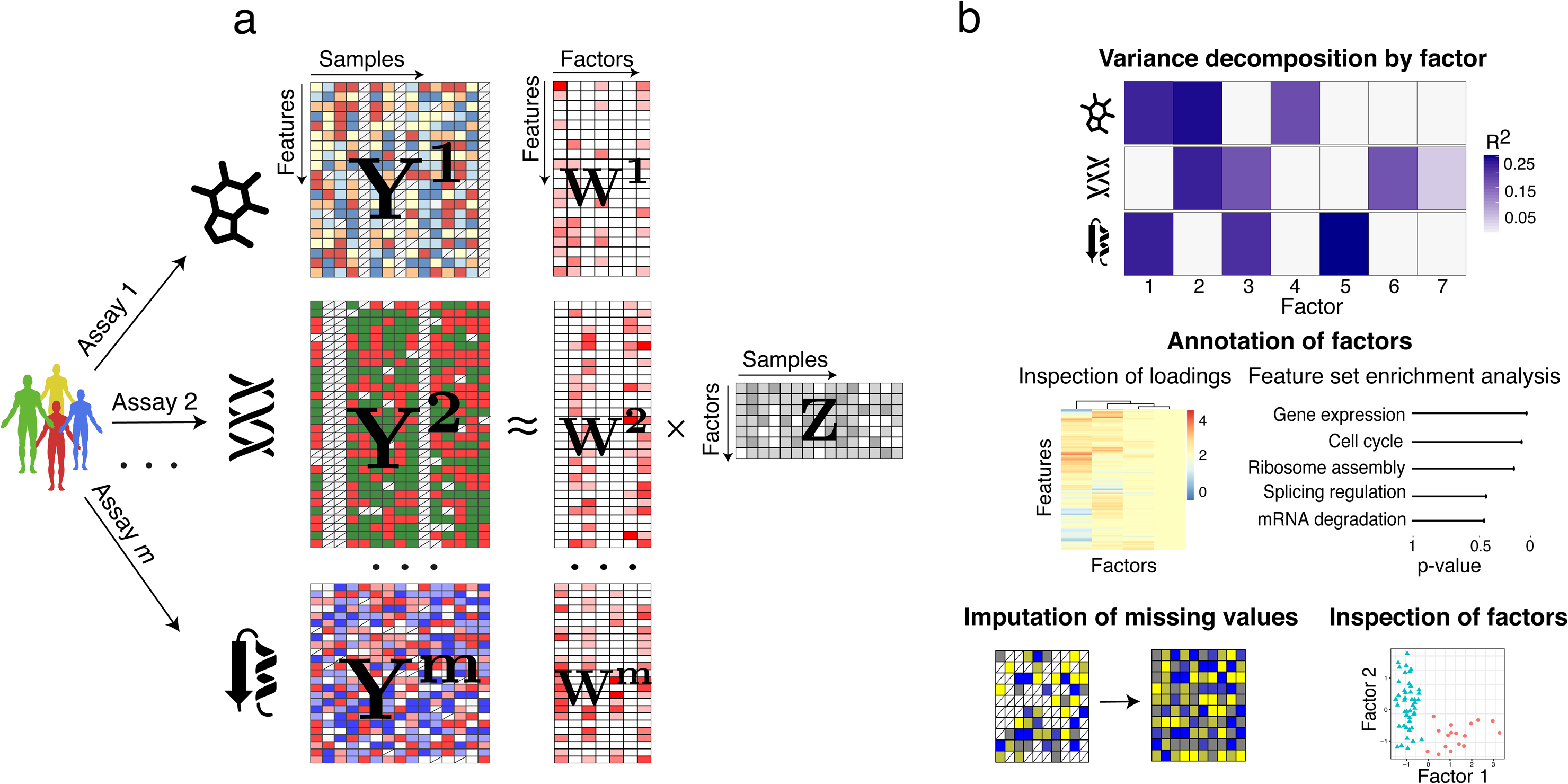
Multi-Omics Factor Analysis: model overview and downstream analyses. **(a)** Model overview: MOFA takes an arbitrary number of M data matrices as input (Y^1^,…, Y^M^), one or more from each data modality, with co-occurrent samples but features that are in general unrelated and that differ in numbers. MOFA decomposes these matrices into a matrix of factors, Z, for each sample and M weight matrices, one for each data modality (loadings W^1^,.., W^M^). White cells in the weight matrices correspond to zeros, i.e. inactive features, whereas the cross symbol in the data matrices denote missing values. (**b**) The fitted MOFA model can be queried for different downstream analyses, including (i) variance decomposition, assessing the proportion of variance explained by each factor in each data modality, (ii) semi-automated factor annotation based on the inspection of loadings and gene set enrichment analysis, (iii) visualization of the samples in the factor space and (iv) imputation of missing values, including missing assays.

Technically, MOFA builds upon the statistical framework of group factor analysis (Bunte et al, 2016; Khan et al, 2014; Klami et al, 2015; Leppäaho & Kaski, 2017; Virtanen et al, 2012; Zhao et al, 2016), which we have adapted to the requirements of multi-omics studies (**Methods**): (i) fast inference based on a variational approximation, (ii) inference of sparse solutions facilitating interpretation, (iii) efficient handling of missing values, and (iv) flexible combination of potentially different likelihood models for each data modality, which enables integrating diverse data types such as binary-, count- and continuous-valued data. The relationship of MOFA to previous approaches (Bunte et al, 2016; Hore et al, 2016; Klami et al, 2015; Leppäaho & Kaski, 2017; Mo et al, 2013; Remes et al, 2015; Shen et al, 2009; Virtanen et al, 2012; Zhao et al, 2016) is discussed in **Methods** and **Appendix Table S3.**

MOFA is implemented as well-documented open-source software that facilitates a range of important downstream analyses, including visualization and automatic characterization of the inferred factors (**Methods**). Taken together, these functionalities provide a powerful and versatile tool for disentangling sources of variation in multi-omic studies.

### Model validation and comparison on simulated data

First, to validate MOFA, we simulated data from its generative model, varying the number and the likelihood model of different views, the number of latent factors and other parameters (**Methods, Appendix Table S1**). We found that MOFA was able to accurately reconstruct the latent dimension, except in settings with large numbers of factors or proportions of missing values (**Appendix Figure S1**). We also found that models with non-Gaussian likelihood models improved the fit when simulating binary or count data (Appendix Figure S2 and S3).

We also compared MOFA to two previously reported latent variable models for multi-omics integration: GFA (Leppäaho & Kaski, 2017) and iCluster (Mo et al, 2013). Over a range of simulations, we observed that GFA and iCluster tended to infer redundant factors (**Appendix Figure S4**) and were less accurate in recovering patterns of factor activity across views (**Appendix Figure S5**). MOFA is also computationally more efficient than GFA and iCluster (**Figure EV1**). For example, the training on the CLL data, which we consider next, required 45 minutes with MOFA vs. 34 hours with GFA and 5-6 days with iCluster.

### Application to Chronic Lymphocytic Leukaemia

We applied MOFA to a study of chronic lymphocytic leukaemia (CLL), which combined ex-vivo drug response measurements with somatic mutation status, transcriptome profiling and DNA methylation assays (Dietrich et al, 2018) (**Fig. 2a**). Notably, nearly 40% of the 200 samples were profiled with some but not all ‘omics types; such a missing value scenario is not uncommon in large cohort studies, and MOFA is designed to cope with it (**Methods; Appendix Figure S1**). MOFA was also configured to combine different likelihood models in order to accommodate the combination of continuous and discrete data types in this study.

**Figure 2:**
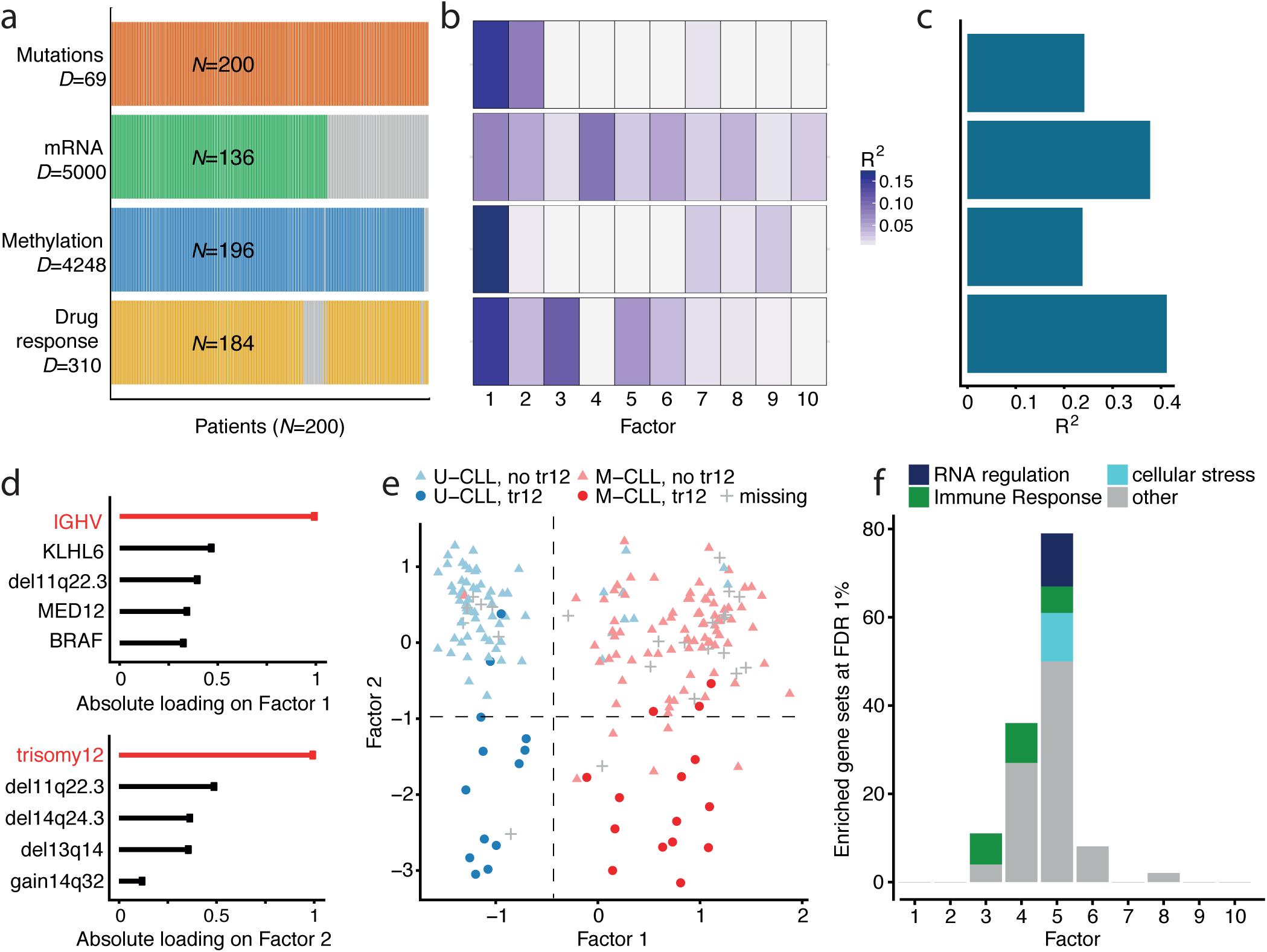
Application of MOFA to a study of chronic lymphocytic leukaemia. **(a)** Study overview and data types. Data modalities are shown in different rows (*D* = number of features) and samples in columns, with missing samples shown using grey bars. **(b)** Proportion of total variance explained (R^2^) by individual factors for each assay and **(c)** cumulative proportion of total variance explained. **(d)** Absolute loadings of the top features of Factors 1 and 2 in the Mutations data. **(e)** Visualisation of samples using Factors 1 and 2. The colors denote the IGHV status of the tumors; symbol shape and color tone indicate chromosome 12 trisomy status. **(f)** Number of enriched Reactome gene sets per factor based on the gene expression data (FDR< 1%). The colors denote categories of related pathways defined as in **Appendix Table S2**.

MOFA identified 10 factors (minimum explained variance 2% in at least one assay; **Methods**). These were robust to algorithm initialisation as well as subsampling of the data (**Appendix Figure S6,7**). The factors were largely orthogonal, capturing independent sources of variation (**Appendix Figure S6**). Among these, Factors 1 and 2 were active in most assays, indicating broad roles in multiple molecular layers (**Fig. 2b**). In contrast, other factors such as Factor 3 or Factor 5 were specific to two data modalities, and Factor 4 was active in a single data modality only. Cumulatively, the 10 factors explained 41% of variation in the drug response data, 38% in the mRNA data, 24% in the DNA methylation data and 24% in the mutation data (**Fig. 2c**).

We also trained MOFA when excluding individual views to probe their redundancy, finding that factors that were active in multiple assays could still be recovered, while the identification of others was dependent on a specific data type (**Appendix Figure S8**). In comparison to GFA (Leppäaho & Kaski, 2017) and iCluster (Mo et al, 2013), MOFA was more consistent in identifying factors across random restarts and their cross-assay activity (**Appendix Figure S9**).

### MOFA identifies important clinical markers in CLL and reveals an underappreciated axis of variation attributed to oxidative stress

As part of the downstream pipeline, MOFA provides different strategies to use the loadings of the features on each factor to identify their etiology (**Fig. 1**). For example, based on the top weights in the mutation data, Factor 1 was aligned with the somatic mutation status of the immunoglobulin heavy-chain variable region gene (IGHV), while Factor 2 aligned with trisomy of chromosome 12 (**Fig. 2d,e**). Thus, MOFA correctly identified two major axes of molecular disease heterogeneity and aligned them with two of the most important clinical markers in CLL (Fabbri & Dalla-Favera, 2016; Zenz et al, 2010) (**Fig. 2d,e**).

IGHV status, the marker corresponding to Factor 1, is a surrogate of the differentiation state of the tumor’s cell of origin and the level of activation of the B-cell receptor. While in clinical practice this axis of variation is generally considered binary (Fabbri & Dalla-Favera, 2016), our results indicate a more complex substructure (**Fig. 3a, Appendix Figure S10**). At the current resolution, this factor was consistent with three subgroup models such as proposed by (Oakes et al, 2016; Queiros et al, 2015) (**Appendix Figure S11**), although there is suggestive evidence for an underlying continuum. MOFA connected this factor to multiple molecular layers (**Appendix Figure S12, S13**), including changes in the expression of genes previously linked to IGHV status (Maloum et al, 2009; Morabito et al, 2015; Plesingerova et al, 2017; Trojani et al, 2012; Vasconcelos et al, 2005) (**Fig. 3b,c**) and with drugs that target kinases in or downstream of the B-cell receptor (**Fig. 3d,e**).

**Figure 3:**
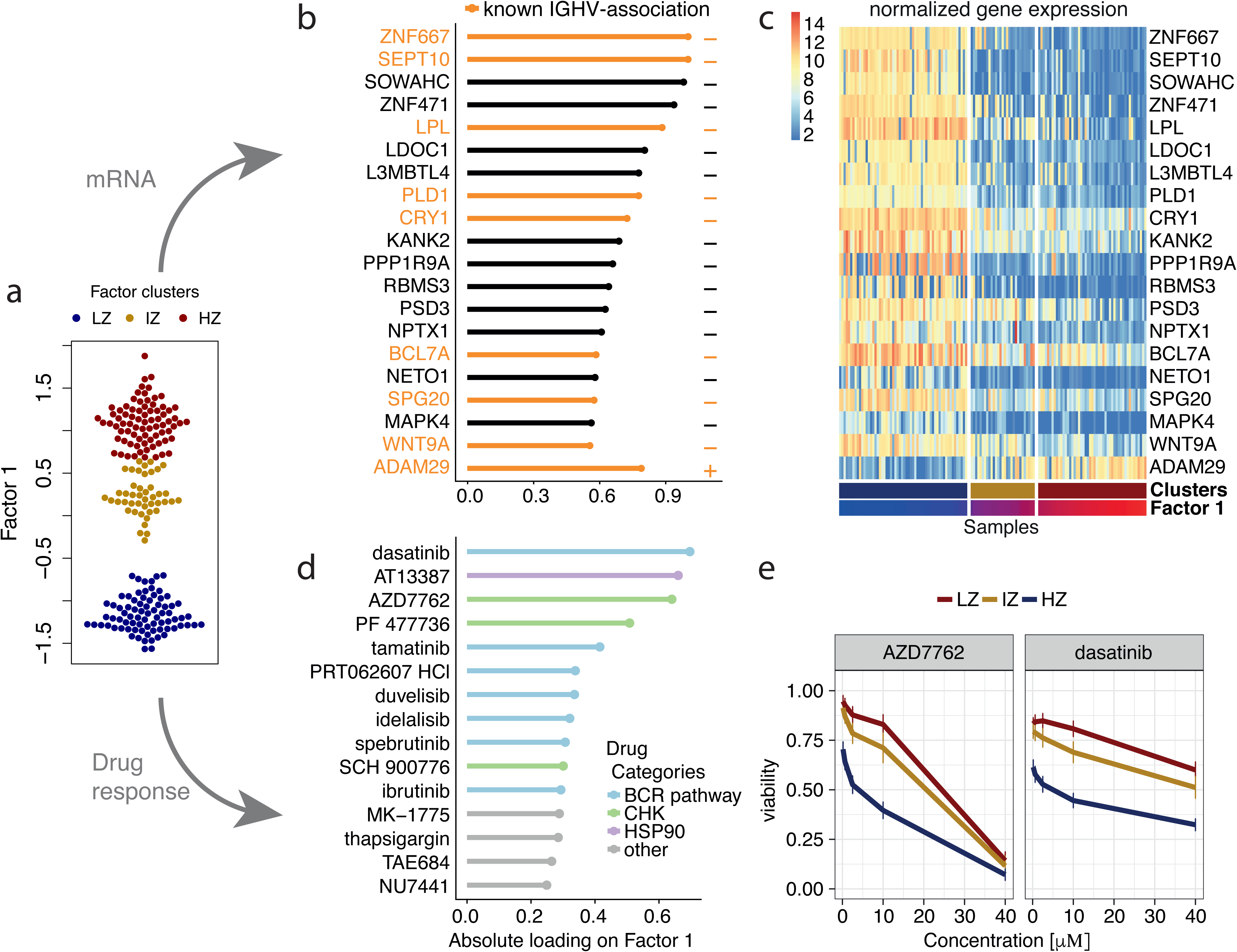
Characterization of the inferred factor associated to the differentiation state of the cell of origin. **(a)** Beeswarm plot with Factor 1 values for each sample with colors corresponding to three clusters found by 3-means clustering with low factor values (LZ), intermediate factor values (IZ) and high factor values (HZ). **(b)** Absolute loadings for the genes with the largest absolute weights in the mRNA data. Plus or minus symbols on the right indicate the sign of the loading. Genes highlighted in orange were previously described as prognostic markers in CLL and associated with IGHV status (Maloum et al, 2009; Morabito et al, 2015; Plesingerova et al, 2017; Trojani et al, 2012; Vasconcelos et al, 2005). **(c)** Heatmap of gene expression values for genes with the largest weights as in **b. (d)** Absolute loadings of the drugs with the largest weights, annotated by target category. **(e)** Drug response curves for two of the drugs, stratified by the clusters as in **a**.

Despite their clinical importance, the IGHV and the trisomy 12 factors accounted for less than 20% of the variance explained by MOFA, suggesting the existence of other sources of heterogeneity. One example is Factor 5, which was active in the mRNA and drug response data. Analysis of the weights in the mRNA revealed that this factor tagged a set of genes enriched for oxidative stress and senescence pathways (**Fig. 2f, Figure EV2a**), with the top weights corresponding to heat shock proteins (HSPs) (**Figure EV2b,c**), genes that are essential for protein folding and are up-regulated upon stress conditions (Åkerfelt et al, 2010; Srivastava, 2002). Although genes in HSP pathways are upregulated in some cancers and have known roles in tumour cell survival (Trachootham et al, 2009), thus far this gene family has received little attention in the context of CLL. Consistent with this annotation based on the mRNA data, we observed that the drugs with the strongest weights on Factor 5 were associated with response to oxidative stress, such as target reactive oxygen species (ROS), DNA damage response and apoptosis (**Figure EV2d,e**).

Factor 4 captured 9% of variation in the mRNA data, and gene set enrichment analysis on the mRNA loadings suggested etiologies related to immune response pathways and T-cell receptor signalling (**Fig. 2f**), likely due to differences in cell type composition between samples: While the samples are comprised mainly of B-cells, Factor 4 revealed a possible contamination with other cell types such as T-cells and monocytes (**Appendix Figure S14**). Factor 3 explained 11% of variation in the drug response data capturing differences in the samples’ general level of drug sensitivity (Geeleher et al, 2016) (**Appendix Figure S15**).

### MOFA identifies outlier samples and accurately imputes missing values

Next, we explored the relationship between inferred factors and clinical annotations, which can be missing, mis-annotated or inaccurate, since they are frequently based on single markers or imperfect surrogates (Westra et al, 2011). Since IGHV status is the major biomarker impacting on clinical care, we assessed the consistency between the inferred continuous Factor 1 and this binary marker. For 176 out of 200 patients, the MOFA factor was in agreement with the clinical IGHV status, and MOFA further allowed for classifying 12 patients that lacked clinically measured IGHV status (**Figure EV3a,b**). Interestingly, MOFA assigned 12 patients to a different group than suggested by their clinical IGHV label. Upon inspection of the underlying molecular data, nine of these cases showed intermediate molecular signatures, suggesting that they are borderline cases that are not well captured by the binary classification; the remaining three cases were clearly discordant (**Figure EV3c,d**). Additional independent drug response assays as well as whole exome sequencing data confirmed that these cases are outliers within their IGHV group (**Fig. EV3e,f**).

As incomplete data is a common problem in studies that combine multiple high-throughput assays, we assessed the ability of MOFA to fill in missing values within assays as well as when entire data modalities are missing for some of the samples. For both imputation tasks, MOFA yielded more accurate predictions than other established imputation strategies, including imputation by feature-wise mean, SoftImpute (Mazumder et al, 2010) and a k-nearest neighbour method (Troyanskaya et al, 2001) (**Figure EV4, Appendix Figure S16**), and MOFA was more robust than GFA, especially in the case of missing assays (**Appendix Figure S17**).

### Latent factors inferred by MOFA are predictive of clinical outcomes

Finally, we explored the utility of the latent factors inferred by MOFA as predictors in models of clinical outcomes. Three of the 10 factors identified by MOFA were significantly associated with time to next treatment (Cox regression, **Methods**, FDR<1%, **Fig. 4a,b**): the cell of origin related Factor 1, and two Factors, 7 and 8, associated with chemo-immunotherapy treatment prior to sample collection. In particular, Factor 7 captures del17p and TP53 mutations as well as differences in methylation patterns of oncogenes (Fluhr et al, 2016; Garg et al, 2014) (**Appendix Figure S18**), while Factor 8 is associated with WNT signalling (**Appendix Figure S19**).

**Figure 4:**
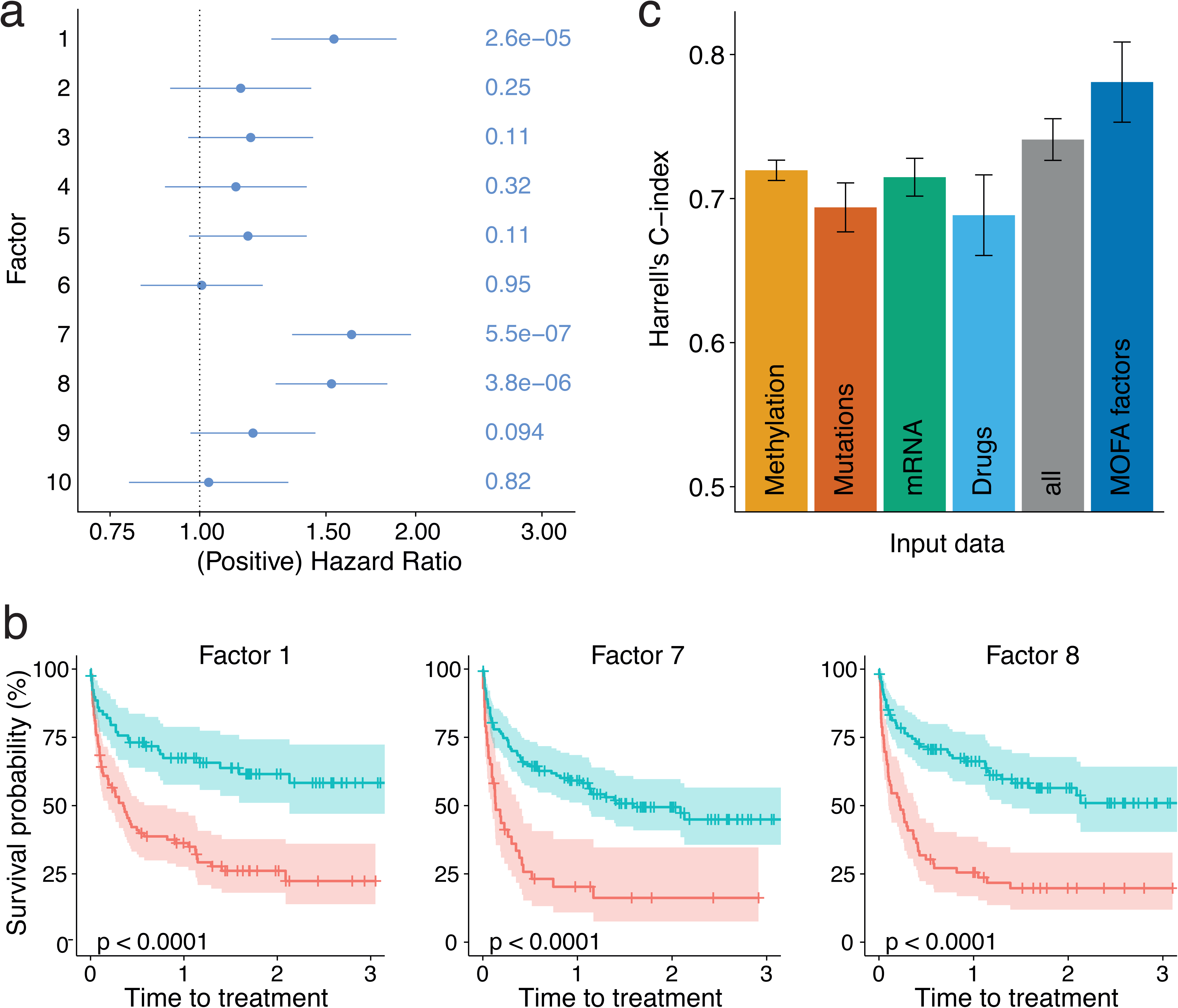
Relationship between clinical data and latent factors. **(a)** Association of MOFA factors to time to next treatment using a univariate Cox models. Error bars denote 95% confidence intervals. Numbers on the right denote p-values for each predictor. **(b)** Kaplan-Meier plots measuring time to next treatment for the individual MOFA factors. The cut-points on each factor were chosen using maximally selected rank statistics(Hothorn & Lausen, 2003), and p-values were calculated using a Log-rank test on the resulting groups. **(c)** Prediction accuracy of time to treatment using multivariate Cox regression trained using the 10 factors derived using MOFA, as well using the first 10 components obtained from PCA applied to the corresponding single data modalities and the full dataset (assessed on hold-out data). Shown are average values of Harrell’s C index from 5-fold cross-validation. Error bars denote standard error of the mean.

We also assessed the prediction performance when combining the 10 MOFA factors in a multivariate Cox regression model. Notably, this model yielded higher prediction accuracy than models using factors derived from conventional PCA (**Fig. 4c**), individual molecular features (**Appendix Figure S20**) or MOFA factors derived from only a subset of the available data modalities (**Appendix Figure S8b,d**) (assessed using cross-validation, **Methods**). Notably, the predictive value of MOFA factors was similar to clinical covariates (such as Lymphocyte doubling time) that are used to guide treatment decisions (**Appendix Figure S21**).

### MOFA reveals coordinated changes in single cells between the transcriptome and the epigenome along a differentiation trajectory

As multi-omic approaches are also beginning to emerge in single cell biology (Angermueller et al, 2016; Clark et al, 2018; Colomé-Tatché & Theis, 2018; Guo et al, 2017; Macaulay et al, 2015), we investigated the potential of MOFA to disentangle the heterogeneity observed in such studies. We applied MOFA to a data set of 87 mouse embryonic stem cells (mESCs), comprising of 16 cells cultured in ‘2i’ media, which induces a naive pluripotency state, and 71 serum-grown cells, which commits cells to a primed pluripotency state poised for cellular differentiation (Angermueller et al, 2016). All cells were profiled using single-cell methylation and transcriptome sequencing, which provides parallel information of these two molecular layers (**Fig. 5a**). We applied MOFA to disentangle the observed heterogeneity in the transcriptome and the CpG methylation at three different genomic contexts: promoters, CpG islands and enhancers.

**Figure 5:**
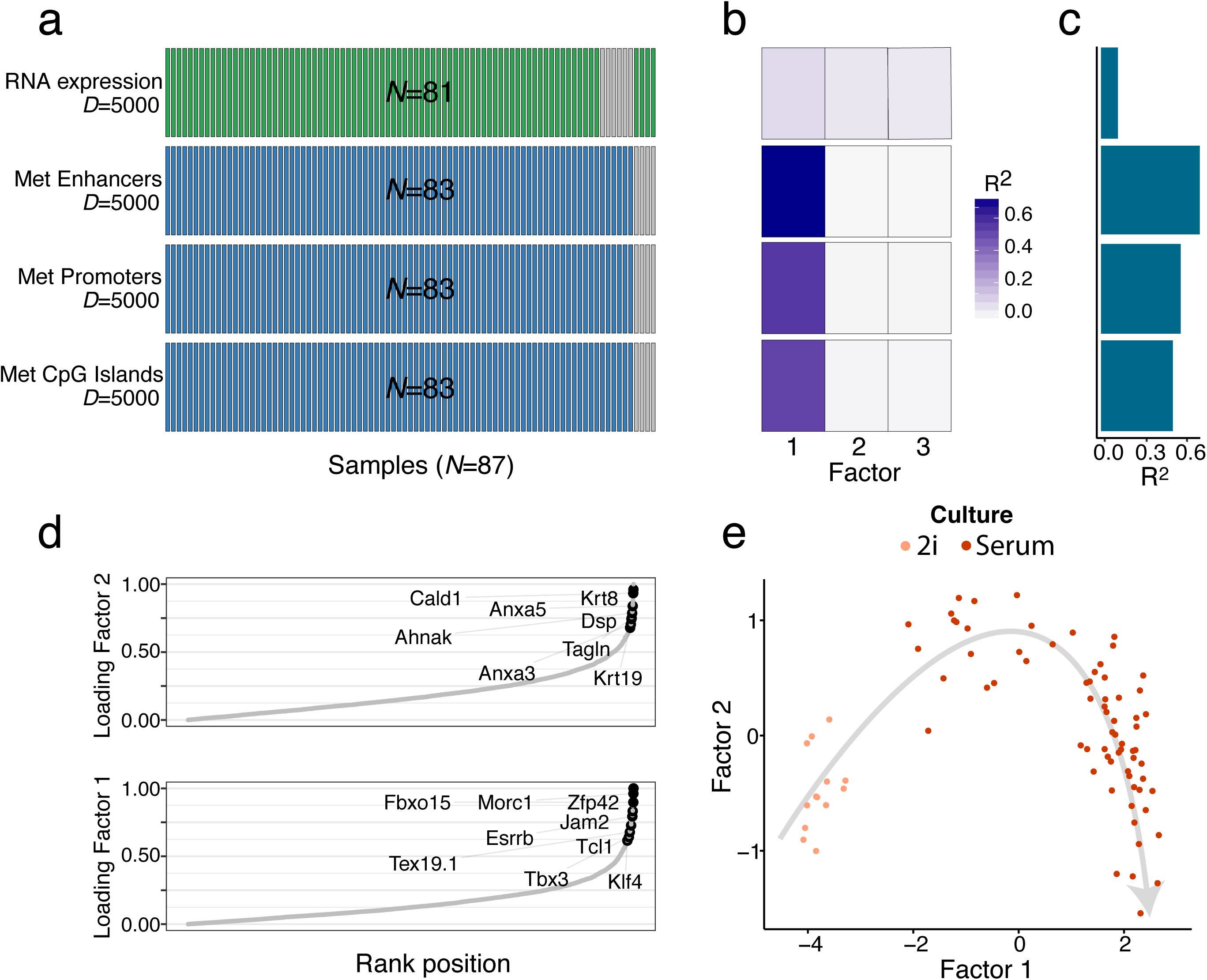
Application of MOFA to a single-cell multi-omics study. **(a)** Study overview and data types. Data modalities are shown in different rows (*D* = number of features) and samples in columns, with missing samples shown using grey bars. **(b)** Fraction of the total variance explained (R^2^) by individual factors for each data modality and **(c)** cumulative proportion of total variance explained. **(d)** Absolute loadings of Factor 1 (bottom) and Factor 2 (top) in the mRNA data. Labeled genes in Factor 1 are known markers of pluripotency (Mohammed et al, 2017) and genes labeled in Factor 2 are known differentiation markers (Fuchs, 1988).**(e)** Scatterplot of Factors 1 and 2. Colors denote culture conditions. The grey arrow illustrates the differentiation trajectory from naive pluripotent cells via primed pluripotent cells to differentiated cells.

MOFA identified 3 factors driving cell-cell heterogeneity (minimum explained variance of 2%, **Methods**): While Factor 1 is shared across all data modalities (7% variance explained in the RNA data and between 53% and 72% in the methylation data sets), Factors 2 and 3 are active primarily in the RNA data (**Fig. 5b,c**). Gene loadings revealed that Factor 1 captured the cell’s transition from naive to primed pluripotent states, pinpointing markers for naive pluripotency such as Rex1/Zpf42, Tbx3, Fbxo15 and Essrb (Mohammed et al, 2017) (**Fig. 5d and Figure EV5a**). MOFA connected these transcriptomic changes to coordinated changes of the genome-wide DNA methylation rate across all genomic contexts (**Figure EV5b**) as previously described both *in vitro*(Angermueller et al, 2016) and *in vivo* (Auclair et al, 2014). Factor 2 captured a second axis of differentiation from the primed pluripotency state to a differentiated state with highest RNA loadings for known differentiation markers such as keratins and annexins (Fuchs, 1988) (**Fig. 5d and Figure EV5c**). Finally, Factor 3 captured the cellular detection rate, a known technical covariate associated with cell quality (Finak et al, 2015) (**Appendix Figure S22**).

Jointly, Factors 1 and 2 captured the entire differentiation trajectory from naive pluripotent cells via primed pluripotent cells to differentiated cells, (**Fig. 5e**), illustrating the importance of learning continuous latent factors rather than discrete sample assignments. Multi-omics clustering algorithms such as SNF (Wang et al, 2014) or iCluster (Mo et al, 2013; Shen et al, 2009) were only capable of distinguishing cellular subpopulations, but not of recovering continuous processes such as cell differentiation (**Appendix Figure S23**).

## Discussion

Multi-Omics Factor Analysis (MOFA) is an unsupervised method for decomposing the sources of heterogeneity in multi-omics data sets. We applied MOFA to high-dimensional and incomplete multi-omics profiles collected from patient-derived tumour samples and to a multi-omics single-cell study of mESCs.

First, in the CLL study, we demonstrated that our method is able to identify major drivers of variation in a clinically and biologically heterogeneous disease. Most notably, our model identified previously known clinical markers as well as novel putative molecular drivers of heterogeneity, some of which were predictive of clinical outcome. Additionally, since MOFA factors capture variations of multiple features and data modalities, inferred factors can help to mitigate assay noise, thereby increasing the sensitivity for identifying molecular signatures compared to using individual features or assays. Our results also demonstrate that MOFA can leverage information from multiple omics layers to accurately impute missing values from sparse profiling datasets and guide the detection of outliers, e.g. due to mislabelled samples or sample swaps.

In a second application, we used MOFA for the analysis of single-cell multi-omics data. This use case illustrates the advantage of learning continuous factors, rather than discrete groups, enabling MOFA to recover a differentiation trajectory by combining information from two sparsely profiled molecular layers.

While applications of factor models for integrating different data types were reported previously (Akavia et al, 2010; Lanckriet et al, 2004; Mo et al, 2013; Shen et al, 2009), MOFA provides unique features (**Methods, Appendix Table S3**) that enable the interpretable reconstruction of the underlying factors and accommodating different data types as well as different patterns of missing data. MOFA is available as open source software and includes semi-automated analysis pipelines allowing for in-depth characterisations of inferred factors. Taken together, this will foster the accessibility of interpretable factor models for a wide range of multi-omics studies.

Although we have addressed important challenges for multi-omics applications, MOFA is not free of limitations. The model is linear, which means that it can miss strongly non-linear relationships between features within and across assays (Buettner & Theis, 2012). Non-linear extensions of MOFA may address this, although as with any models in high-dimensional spaces, there will be trade-offs between model complexity, computational efficiency and interpretability (Damianou et al, 2016). A related area of work is to incorporate prior information on the relationships between individual features. For example, future extensions could make use of pathway databases within each omic type (Buettner et al, 2017) or priors that reflect relationships given by the ‘dogma of molecular biology’. In addition, new likelihoods and noise models could expand the value of MOFA in data sets with specific statistical properties that hamper the application of traditional statistical methods, including zero-inflated data (i.e. scRNA-seq (Pierson & Yau, 2015)) or binomial distributed data (i.e. splicing events (Huang & Sanguinetti, 2017)). Finally, while here we use approximate Bayesian inference and focus attention on the resulting point estimates of inferred factors, future extensions could attempt a more comprehensive Bayesian treatment that propagates evidence strength and estimation uncertainties to diagnostics and downstream analyses.

## Methods

### Code availability

An open source implementation of MOFA is available from https://github.com/bioFAM/MOFA. Code to reproduce all the analyses presented is available at https://github.com/PMBio/MOFA_CLL.

### Data availability

The CLL data were obtained from (Dietrich et al, 2018) and are available at the European Genome-phenome Archive under accession EGAS00001001746 and data tables as R objects can be downloaded from http://pace.embl.de/. The single-cell data were obtained from (Angermueller et al, 2016) and are available in the Gene Expression Omnibus under accession GSE74535. All data used are contained within the MOFA vignettes and can be downloaded as from https://github.com/bioFAM/MOFA.

### Multi-Omic Factor Analysis Model

Starting from M data matrices **Y**^1^,..,**Y**^*M*^ of dimensions *N*× *D_m_*, where N is the number of samples and *D_m_* the number of features in data matrix m, MOFA decomposes these matrices as

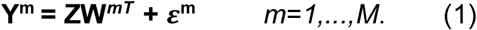

Here, **Z** denotes the factor matrix (common for all data matrices) and **W**^*m*^ denote the weight matrices for each data matrix *m* (also referred to as view *m* in the following). *ε*^m^ denotes the view-specific residual noise term, with its form depending on the specifics of the data-type (see section *Noise model*).

The model is formulated in a probabilistic Bayesian framework, where we place prior distributions on all unobserved variables of the model (see plate diagram in **Appendix Figure S24**), i.e. the factors **Z**, the weight matrices **W**^*m*^ and the parameters of the residual noise term. In particular, we use a standard normal prior for the factors **Z** and employ sparsity priors for the weight matrices (see next section).

### Model regularization

An appropriate regularization of the weight matrices is essential for the model’s ability to disentangle variation across data sets and yield interpretable factors. MOFA uses a two-level regularization: The first level encourages view- and factor-wise sparsity, thereby allowing to directly identify which factor is active in which view. The second level encourages feature-wise sparsity, thereby typically resulting in a small number of features with active weights. To encode these sparsity levels we combine an Automatic Relevance Determination prior for the first type of the sparsity with a spike-and-slab prior for the second. For amenable inference we model the spike-and-slab prior by parameterizing the weights as a product of a Bernoulli distributed random variable and a normally distributed random variable: 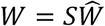 where 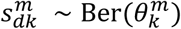 and 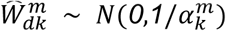. To automatically learn the appropriate level of regularization for each factor and view, we use uninformative conjugate prior on 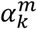 which controls the strength of factor *k* in view *m*, and on 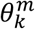 which determines the feature-wise sparsity level of factor *k* in view *m* (**see Appendix Supplementary Methods, section 2** for details).

### Noise model

MOFA supports the combination of different noise models to integrate diverse data types, including continuous, binary and count data. A standard noise model for continuous data is the Gaussian noise model assuming iid heteroscedastic residuals *ε^m^* i.e. 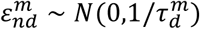, with Gamma prior on the precision parameters 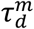 MOFA further supports noise models for binary and count data that are not appropriately modelled using a Gaussian likelihood. In the current version, MOFA models count data using a Poisson model and binary data by using a Bernoulli model. Here, the model likelihood is given by 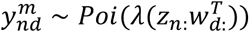 and 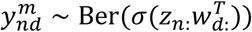 respectively, where *λ*(*x*) = log(1 + *e^x^*) and *σ* denotes the logistic function *σ*(*x*) = (1 + *e^-x^*)^-1^.

### Parameter inference

For scalability, we make use of a variational framework with a mean-field approximation (Blei et al, 2017). The key idea is to approximate the intractable posterior distribution from a simpler class of distributions by minimizing the Kullback-Leibler divergence to the posterior, or equivalently, maximizing the evidence lower bound (ELBO). Convergence of the algorithm can be monitored based on the ELBO. A short introduction to variational inference and details on the algorithm for MOFA can be found in **Appendix Supplementary Methods, section 3.** To enable an efficient inference for non-Gaussian likelihoods we employ variational lower bounds on the likelihood (Jaakkola & Jordan, 2000; Seeger & Bouchard, 2012) (see **Appendix Supplementary Methods, section 4**).

### Model training and selection

An important part of the training is the determination of the number of factors. Factors are automatically inactivated by the model with help of the ARD prior as described in *Model regularization*. In practice, factors are pruned during training using a minimum fraction of variance explained threshold that needs to be specified by the user. Alternatively, the user can fix the number of factors and the minimum variance criterion is ignored. In the analyses presented we initialised the models with *K*=25 factors and they were pruned during training using a threshold of variance explained of 2%. For details on the implementation as well as practical considerations for training and choice of the threshold parameter refer to **Appendix Supplementary Methods, section 5**.

While the inferred factors are robust under different initializations (e.g. **Appendix Figure S6c,d**) the optimization landscape is non-convex and the algorithm is not guaranteed to identify a global optimum. Results presented here are based on 10-25 random restarts, selecting the model with the highest ELBO (e.g. **Appendix Figure S6b**).

### Downstream analysis for factor interpretation and annotation

As part of MOFA we provide the R package *MOFAtools*, containing a semi-automated pipeline for the characterisation and interpretation of the latent factors. In all downstream analyses we use the expectations of the model components under the posterior distributions inferred by the variational framework.

The first step, after a model has been trained, is to disentangle the variation explained by each factor in each view. To this end, we compute the fraction of total variance explained (R^2^) by factor *k* in view *m* as

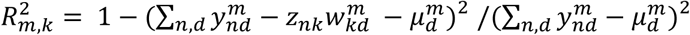

as well as the fraction of variance explained per view taking into account all factors

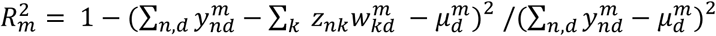

Here, 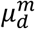 denotes the feature-wise mean. Subsequently, each factor is characterised by three complementary analyses:

1. **Ordination of the samples in factor space**: Visualise a low-dimensional representation of the main drivers of sample heterogeneity.
2. **Inspection of top features with largest weight**: The loadings can give insights into the biological process underlying the heterogeneity captured by a latent factor. Due to scale differences between assays, the weights of different views are not directly comparable. For simplicity, we scale each weight vector by its absolute value.
3. **Feature set enrichment analysis**: We combine the signal from functionally related sets of features (e.g., gene sets) to derive a feature-set based annotation. By default, we use a parametric t-test comparing the means of the foreground set (the weights of features that belong to a set G) and the background set (the weights of features that do not belong to the set G), similar to (Frost et al, 2015).

### Relationship to existing methods

MOFA builds upon the statistical framework of group factor analysis (Bunte et al, 2016; Khan et al, 2014; Klami et al, 2015; Leppäaho & Kaski, 2017; Virtanen et al, 2012; Zhao et al, 2016) and shares components of the iCluster methods (Mo et al, 2013; Shen et al, 2009) as shown in **Appendix Table S3**. Here we describe the connections in more detail:

#### iCluster

In contrast to MOFA, iCluster uses in a each view the same extent of regularization for all factors, which may be sufficient for the purpose of clustering (the primary application of iCluster), however it results in a reduced ability for distinguishing factors that drive variation in distinct subsets of views (**Appendix Figure S5**). Additionally, unlike MOFA and GFA, iCluster does not handle missing values and is computationally demanding (**Figure EV1**), as it requires re-fitting the model for a large range of different penalty parameters and choices of the model dimension.

#### Group factor analysis

While the underlying model of MOFA is closely connect to the most recent GFA implementation (Leppäaho & Kaski, 2017), GFA is restricted to Gaussian observation noise. In terms of implementation, MOFA adds a burn-in period during training without sparsity constraints to avoid early splitting of factors and actively drops factors below the variance threshold as described in *Model training and selection*. In contrast, GFA directly uses sparsity constraints from the beginning and maintains all factors that have non-zero weights. In terms of inference, MOFA is implemented using a variational approach while GFA uses a Gibbs sampling scheme. In terms of scalability (**Figure EV1**), both methods are linear in the model’s parameters. The higher intercept and slope for GFA is mainly driven by the presence of missing values in the data. This, together with the inability to drop factors (**Appendix Figure S4**) renders GFA considerably slower in applications to real data.

### Details on the simulation studies

#### Model validation

To validate MOFA we simulated data from the generative model for a varying number of views (M=1,3,…,21), features (D=100,500,…,10000), factors (K=5,10,…,60), missing values (from 0% to 90%) as well as for non-Gaussian likelihoods (Poisson, Bernoulli) (see **Appendix Table S1** for simulation parameters). We assessed the ability of MOFA to recover the true number of factors in the different settings across 10 realizations for every configuration. All trials were started with a high number of factors (*K*=100), and inactive factors were pruned as described in the *Model training and selection* section.

#### Model comparison

To compare MOFA with other approaches we simulated data from the generative model with *K*_true_=10 factors, *M*=3 views, *N*=100 samples, *D*=5,000 features each and 5% missing values (missing at random). For each of the three views we used a different likelihood model: continuous data was simulated with a Gaussian distribution, binary data with a Bernoulli distribution and count data with a Poisson distribution. Except for the non-Gaussian likelihood extension, both methods share the same underlying generative model, which ensures a fair comparison. We fit ten realization of the MOFA and GFA models starting with *K*_initial_=20 factors, and let the method learn the true number of factors. To assess scalability, the same simulation setting was used varying one of the simulation parameters at a time (number of factors *K*, number of features *D*, number of samples *N* and number of views *M*, all Gaussian). To compare the ability to reconstruct factor activity patterns we simulated data from the generative model for *K*_true_=10 and *K*_true_=15 factors (*M*, *N*, *D* as before, no missing values, only Gaussian views), where factors were set to either active or inactive in a specific view by sampling the parameter 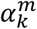 from {1,10^3^}. **Appendix Table S1** shows in more detail the simulation parameters used in each setting.

### Details on the CLL analysis

#### Data processing

The data was obtained from (Dietrich et al, 2018) where details on the data generation and processing can be found. Briefly, the data consist of somatic mutations (combination of targeted and whole exome sequencing), RNA expression (RNA-Seq), DNA methylation (Illumina arrays) and *ex-vivo* drug response screens (ATP-based CellTiter Glo assay). For the training of MOFA we included 62 drug response measurements (excluding NSC 74859 and bortezomib due to bad quality) at five concentrations each (*D*=310) with a threshold at 1.1 to remove outliers. Mutations were considered if present in at least 3 samples (*D*=69). Low counts from RNAseq data were filtered out and the data was normalized using the *estimateSizeFactors* and *varianceStabilizingTransformation* function of DESeq2 (Love et al, 2014). For training we used the top *D*=5000 most variable mRNAs after exclusion of genes from the Y chromosome. Methylation data was transformed to M-values and we extracted the top 1% most variable CpG sites excluding sex chromosomes (*D*=4248). We included patients diagnosed with CLL and having data in at least two views into the MOFA model leading to a total of *N*=200 samples.

#### Model training and selection

We trained MOFA on the data for 25 random initializations with a variance threshold of 2% and selected a model for downstream analysis as described in *Model training and selection*.

#### Gene set enrichment analysis

Gene set enrichment analysis was performed on the Reactome gene sets (Fabregat et al, 2015) as described above. Resulting p-values are adjusted for multiple testing on each factor using Benjamini-Hochberg (Benjamini & Hochberg, 1995) procedure to control the false discovery rate at 1%.

#### Imputation

To compare imputation performance, we trained MOFA on the subset of samples with all measurements (*N*=121) and masked at random either single values or all measurements for given samples in the drug response. After model training the masked values were imputed directly from the model equation (1) and the accuracy was assessed in terms of mean squared error on the true (masked) values. For both settings we fixed the number of factors in MOFA to *K*=10. To investigate the dependence on *K* for imputation and compare to GFA we re-ran the same masking experiments varying *K*=1,…,20.

#### Survival Analysis

Associations of the inferred factors to clinical outcome were assessed using patients’ time to next treatment as response variable in a Cox model on all samples having this information, i.e. *N*=174 of which 96 are uncensored cases. For univariate associations (as shown in **Figure 4a, Appendix Figure S21**) we scaled all predictors to ensure comparability of the hazard ratios and we rotated factors, which are rotational invariant, such that their Hazard ratio is greater or equal to 1. To investigate the predictive power of different datasets, we used a multivariate Cox model and compared Harrell's C-index of predictions in a stratified 5-fold cross-validation scheme. As predictors we included the top 10 principal components on the data of each single view, a concatenated data set ('all') as well as the ten MOFA factors. Missing values in a view were imputed by the feature-wise mean. In a second set of models we used the complete set of all features in a view with a ridge penalty in the Cox model as implemented in the R package *glmnet*. For the Kaplan-Meier plots an optimal cut-point on each factor was determined to define the two groups using the maximally selected rank statistics as implemented in the R package *survminer*with p-values based on a Log-Rank test between the resulting groups.

### Details on the scMT analysis

The data were obtained from (Angermueller et al, 2016), where details on the data generation and pre-processing can be found. Briefly for each CpG site we calculated a binary methylation rate from the ratio of methylated read counts to total read counts. RNA expression data were normalised using (Lun et al, 2016). To fit MOFA, we considered the top 5000 most variable genes with a maximum dropout of 90%, and the top 5000 most variable CpG sites with a minimum coverage of 10% across cells. Model selection was performed as described for the CLL data and factors were inactivated below a minimum explained variance of 2%. For the clustering analysis using SNF and iCluster, the optimal number of clusters was selected using BIC criterion.

## Acknowledgements

We thank everybody involved in the generation and analysis of the original CLL study for sharing their data and analysis ahead of publication, especially M. Oleś for providing the associated data package and to J. Lu, J. Hüllein and A. Mock for discussions on CLL biology. The work was supported by the European Union (Horizon 2020 project SOUND and project PanCanRisk).

## Author contributions

RA and BV contributed equally and are listed alphabetically. FB, DA and OS conceived the model.

RA, DA and BV implemented the model.

TZ, SD, WH designed the CLL study and generated the data. RA and BV performed the analysis.

RA, BV, DA, TZ, SD, WH, OS, FB, JM interpreted the results. RA, BV, OS, WH, FB conceived the project.

RA, BV, OS, FB, WH wrote the manuscript.

OS, WH, FB, JM supervised the project.

---

**Expanded View Figure 1.**
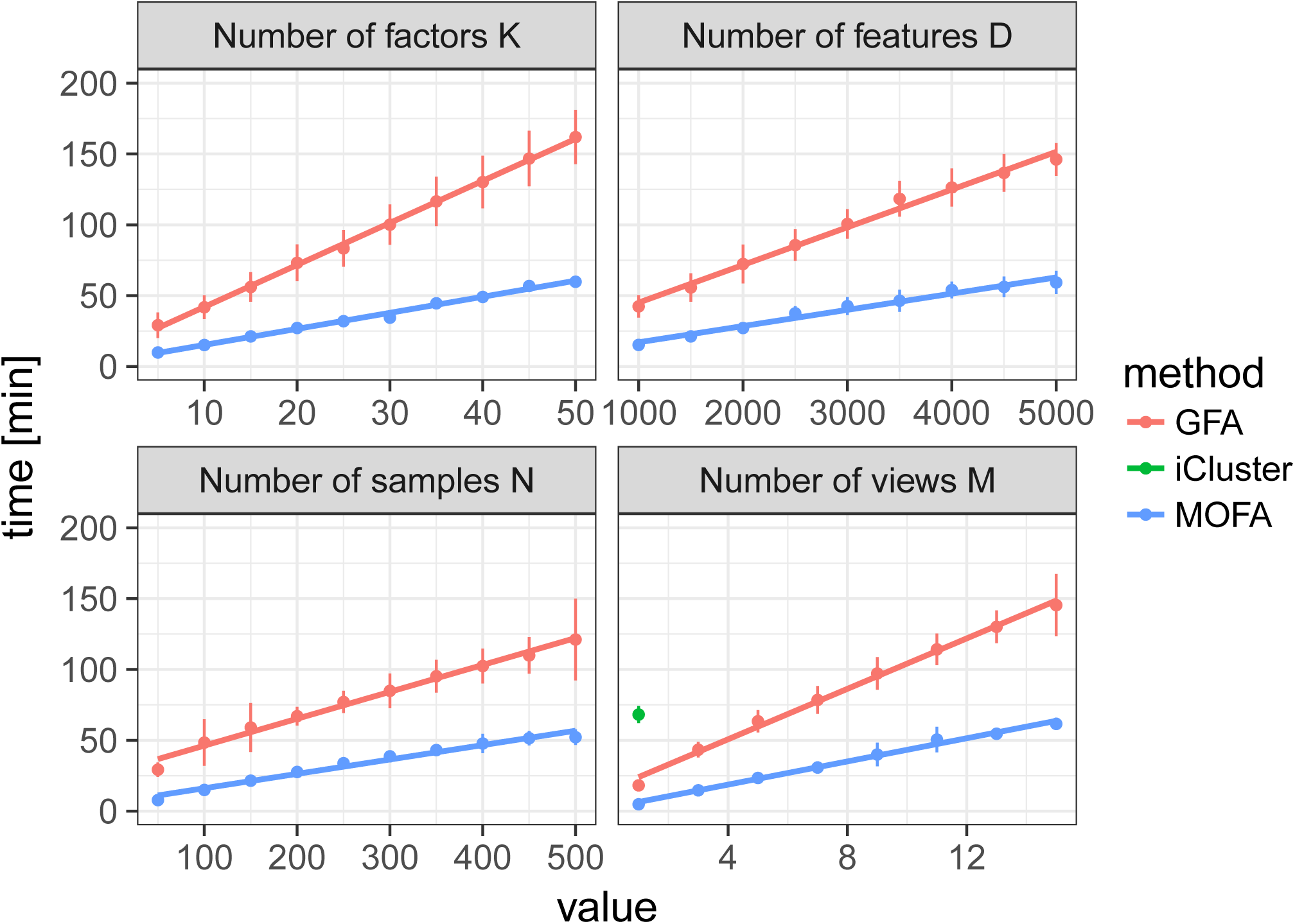
Scalability of MOFA, GFA and iCluster. Time required for model training for GFA (red), MOFA (blue) and iCluster (green) as a function of number of factors *K*, number of features D, number of samples *N* and number of views M. Baseline parameters were *M=3, K=10, D=1000* and *N=100* and 5% missing values. Shown are average time across 10 trials, error bars denote standard deviation. iCluster is only shown for the lowest *M* as all other settings require on average more than 200 minutes for training.

**Expanded View Figure 2.**
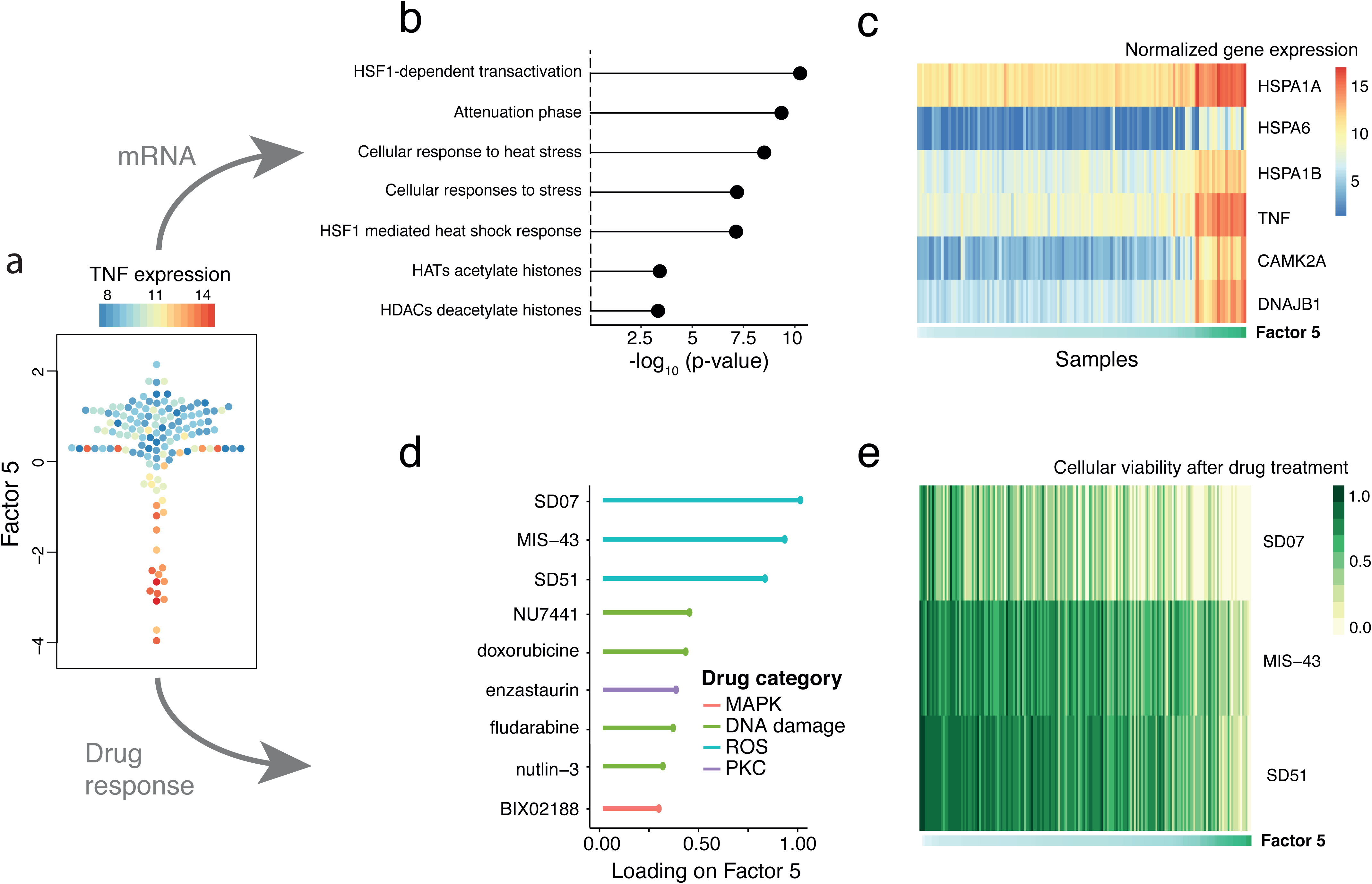
Characterization of Factor 5 (oxidative stress response factor) in the CLL data. **(a)** Beeswarm plot of Factor 5. Colors denote the expression of TNF, an inflammatory stress marker. **(b)** Gene set enrichment for the top Reactome pathways in the mRNA data (t-test, **Methods). (c)** Heatmap of gene expression values for the six genes with largest loading. Samples are ordered by their factor values. **(d)** Scaled loadings for the top drugs with the largest loading, annotated by target category. **(e)** Heatmap of drug response values for the top three drugs with largest loading.

**Expanded View Figure 3.**
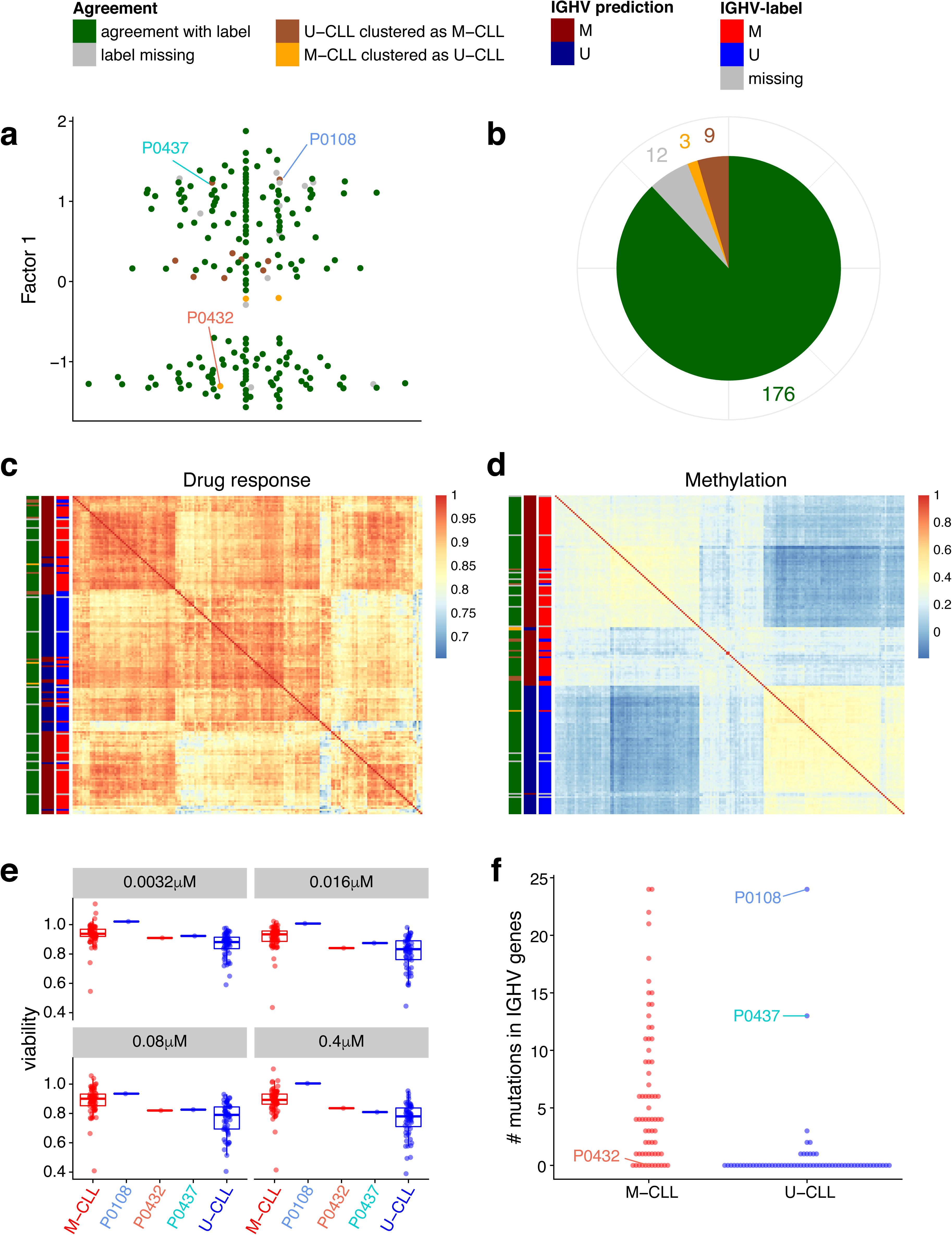
Prediction of IGHV status based on Factor 1 in the CLL data and validation on outlier cases on independent assays. **(a)** Beeswarm plot of Factor 1 with colours denoting agreement between predicted and clinical labels as in **(b). (b)** Pie chart showing total numbers for agreement of imputed labels with clinical label. **(c)** Sample-to-sample correlation matrix based on drug response data. **(d)** Sample-to-sample correlation matrix based on methylation data. **(e)** Drug response to ONO-4509 (not included in the training data): Boxplots for the viability values in response to ONO-4509. The three outlier samples are shown in the middle, on the left and right the viabilities of the other M-CLL and U-CLL samples are shown, respectively. The panels show different drug concentrations tested. **(b)** Whole exome sequencing data on IGHV genes (not included in the training data): the number of mutations found on IGHV genes using whole exome sequencing is shown on the y-axis, separately for U-CLL and M-CLL samples. The three outlier samples are labelled.

**Expanded View Figure 4.**
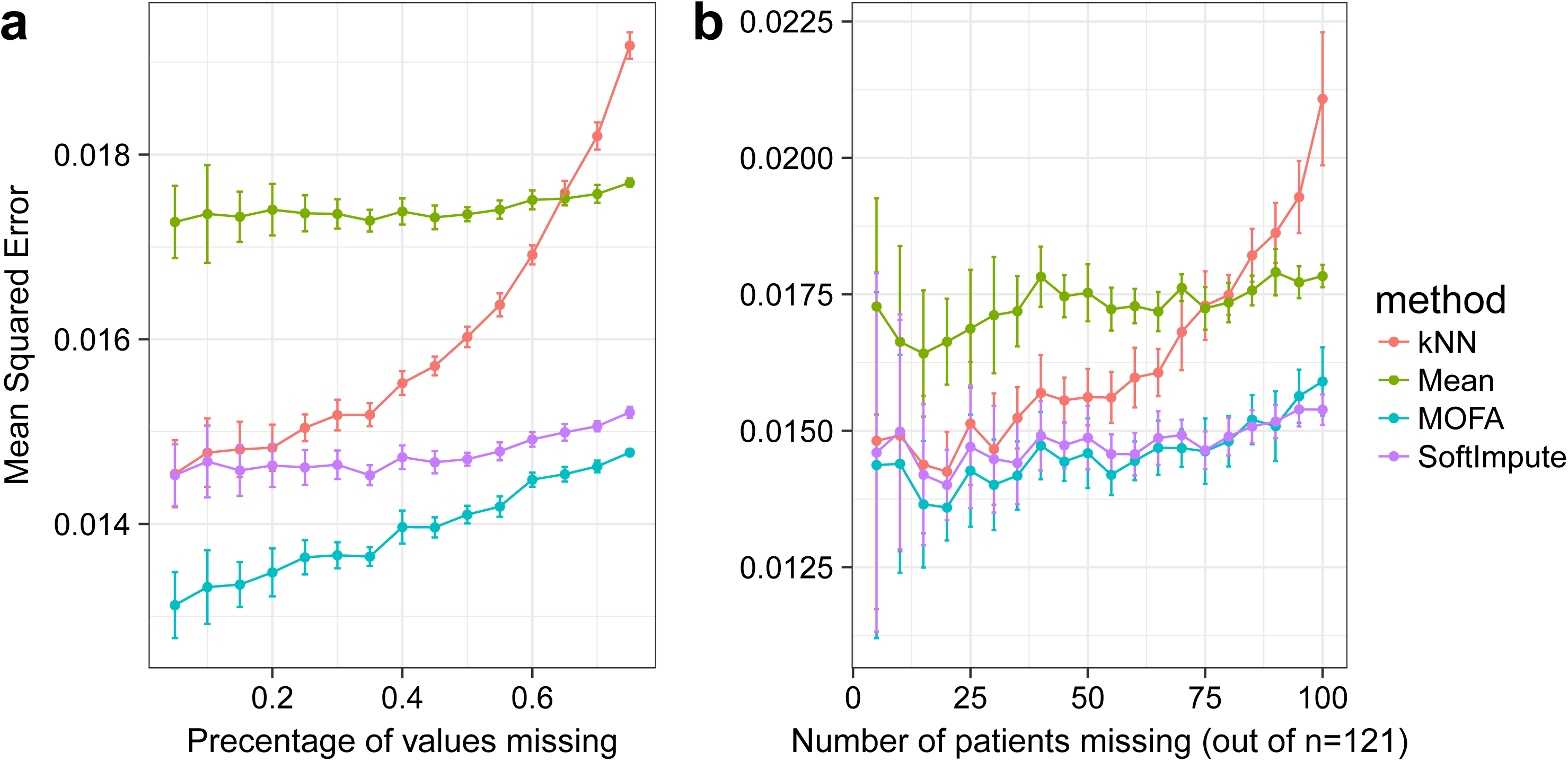
Imputation of missing values in the drug response assay of the CLL data. Considered were MOFA, SoftImpute, imputation by feature-wise mean (Mean) and k-nearest neighbour (kNN). Shown are averages of the mean squared error (MSE) across 15 imputation experiments for increasing fractions of missing data, considering **(a)** values missing at random and **(b)** entire assay missing for samples at random. Error bars denote plus or minus two standard error.

**Expanded View Figure 5.**
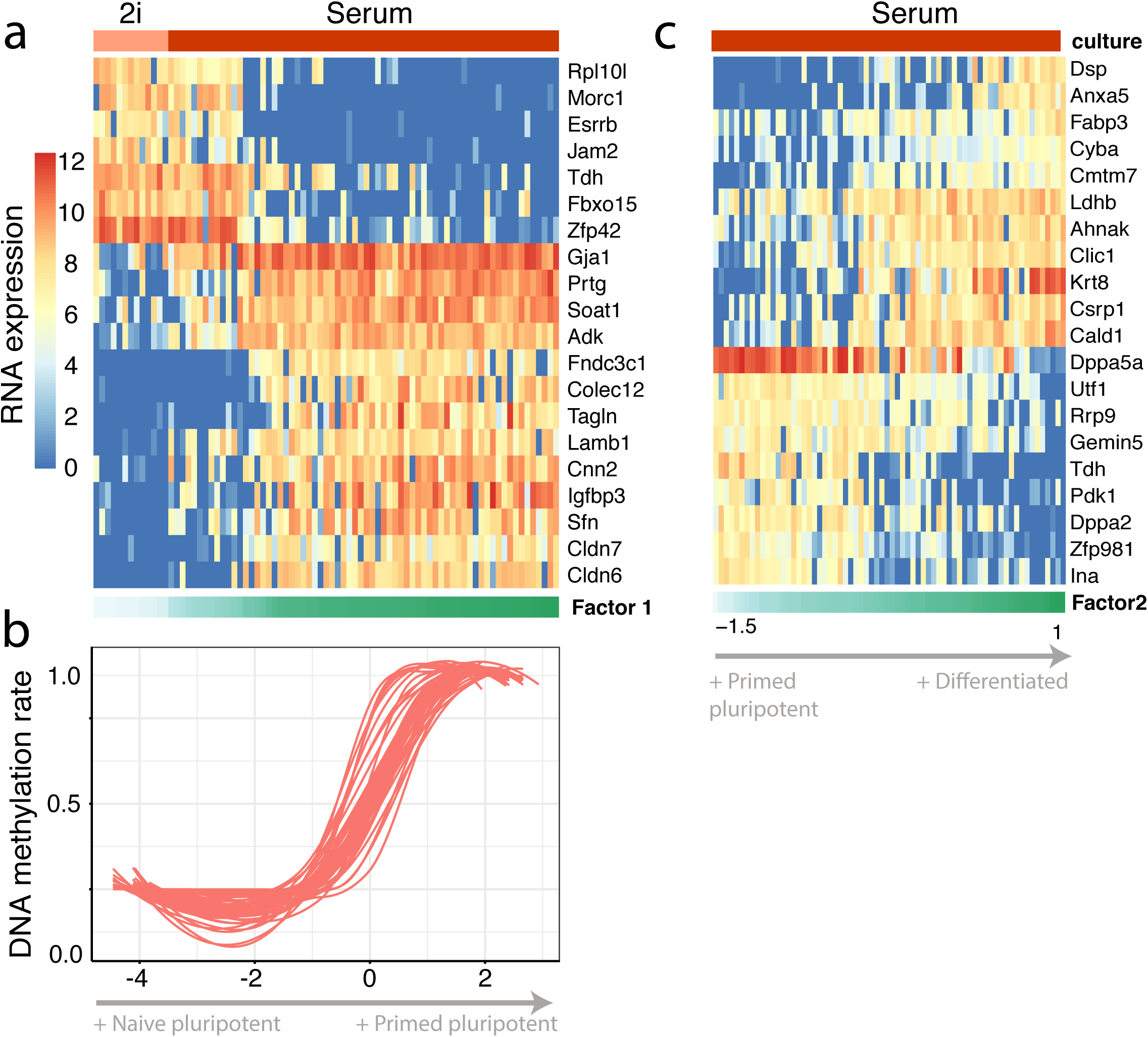
Transcriptomic and epigenetic changes associated with Factor 1 in the scMT data. **(a)** RNA expression changes for the top 20 genes with largest weight on Factor 1. **(b)** DNA methylation rate changes for the top 20 CpG sites with largest weight. Shown is a non-linear loess regression model fit per CpG site. **(c)** RNA expression changes for the top 20 genes with largest weight on Factor 2.

## References

Akavia UD, Litvin O, Kim J, Sanchez-Garcia F, Kotliar D, Causton HC, Pochanard P, Mozes E, Garraway LA, Pe’er D (2010) An integrated approach to uncover drivers of cancer. Cell 143: 10051017

Åkerfelt M, Morimoto RI, Sistonen L (2010) Heat shock factors: integrators of cell stress, development and lifespan. Nature reviews Molecular cell biology 11: 545

Angermueller C, Clark SJ, Lee HJ, Macaulay IC, Teng MJ, Hu TX, Krueger F, Smallwood S, Ponting CP, Voet T, Kelsey G, Stegle O, Reik W (2016) Parallel single-cell sequencing links transcriptional and epigenetic heterogeneity. Nature methods 13: 229–232

Auclair G, Guibert S, Bender A, Weber M (2014) Ontogeny of CpG island methylation and specificity of DNMT3 methyltransferases during embryonic development in the mouse. Genome biology 15: 545

Benjamini Y, Hochberg Y (1995) Controlling the false discovery rate: a practical and powerful approach to multiple testing. Journal of the royal statistical society Series B (Methodological): 289–300

Blei DM, Kucukelbir A, McAuliffe JD (2017) Variational inference: A review for statisticians. Journal of the American Statistical Association 112: 859–877

Buettner F, Pratanwanich N, McCarthy DJ, Marioni JC, Stegle O (2017) f-scLVM: scalable and versatile factor analysis for single-cell RNA-seq. Genome biology 18: 212

Buettner F, Theis FJ (2012) A novel approach for resolving differences in single-cell gene expression patterns from zygote to blastocyst. Bioinformatics 28: i626–i632

Bunte K, Leppaaho E, Saarinen I, Kaski S (2016) Sparse group factor analysis for biclustering of multiple data sources. Bioinformatics 32: 2457–2463

Cancer Genome Atlas Research Network (2017) Comprehensive and Integrative Genomic Characterization of Hepatocellular Carcinoma. Cell 169: 1327–1341 e1323

Chen L, Ge B, Casale FP, Vasquez L, Kwan T, Garrido-Martín D, Watt S, Yan Y, Kundu K, Ecker S (2016) Genetic drivers of epigenetic and transcriptional variation in human immune cells. Cell 167: 1398–1414. e1324

Clark SJ, Argelaguet R, Kapourani C-A, Stubbs TM, Lee HJ, Alda-Catalinas C, Krueger F, Sanguinetti G, Kelsey G, Marioni JC (2018) scNMT-seq enables joint profiling of chromatin accessibility DNA methylation and transcription in single cells. Nature communications 9: 781

Colomé-Tatché M, Theis F (2018) Statistical single cell multi-omics integration. Current Opinion in Systems Biology 7: 54–59

Consortium G (2015) The Genotype-Tissue Expression (GTEx) pilot analysis: Multitissue gene regulation in humans. Science 348: 648–660

Damianou A, Lawrence ND, Ek CH (2016) Multi-view Learning as a Nonparametric Nonlinear Inter-Battery Factor Analysis. arXiv preprint arXiv:160404939

Dietrich S, Oleś M, Lu J, Sellner L, Anders S, Velten B, Wu B, Hüllein J, da Silva Liberio M, Walther T (2018) Drug-perturbation-based stratification of blood cancer. The Journal of clinical investigation 128: 427–445

Fabbri G, Dalla-Favera R (2016) The molecular pathogenesis of chronic lymphocytic leukaemia. Nature reviews Cancer 16: 145–162

Fabregat A, Sidiropoulos K, Garapati P, Gillespie M, Hausmann K, Haw R, Jassal B, Jupe S, Korninger F, McKay S (2015) The reactome pathway knowledgebase. Nucleic acids research 44:D481–D487

Finak G, McDavid A, Yajima M, Deng J, Gersuk V, Shalek AK, Slichter CK, Miller HW, McElrath MJ, Prlic M (2015) MAST: a flexible statistical framework for assessing transcriptional changes and characterizing heterogeneity in single-cell RNA sequencing data. Genome biology 16: 278

Fluhr S, Boerries M, Busch H, Symeonidi A, Witte T, Lipka DB, Mücke O, Nöllke P, Krombholz CF, Niemeyer CM (2016) CREBBP is a target of epigenetic, but not genetic, modification in juvenile myelomonocytic leukemia. Clinical epigenetics 8: 50

Frost HR, Li Z, Moore JH (2015) Principal component gene set enrichment (PCGSE). BioData mining 8: 25

Fuchs E (1988) Keratins as biochemical markers of epithelial differentiation. Trends in Genetics 4:277–281

Garg R, Benedetti LG, Abera MB, Wang H, Abba M, Kazanietz MG (2014) Protein kinase C and cancer: what we know and what we do not. Oncogene 33: 5225–5237

Geeleher P, Cox NJ, Huang RS (2016) Cancer biomarker discovery is improved by accounting for variability in general levels of drug sensitivity in pre-clinical models. Genome biology 17: 190

Gerstung M, Pellagatti A, Malcovati L, Giagounidis A, Porta MG, Jadersten M, Dolatshad H, Verma A, Cross NC, Vyas P, Killick S, Hellstrom-Lindberg E, Cazzola M, Papaemmanuil E, Campbell PJ, Boultwood J (2015) Combining gene mutation with gene expression data improves outcome prediction in myelodysplastic syndromes. Nature communications 6: 5901

Guo F, Li L, Li J, Wu X, Hu B, Zhu P, Wen L, Tang F (2017) Single-cell multi-omics sequencing of mouse early embryos and embryonic stem cells. Cell research 27: 967–988

Hasin Y, Seldin M, Lusis A (2017) Multi-omics approaches to disease. Genome biology 18: 83

Hore V, Viñuela A, Buil A, Knight J, McCarthy MI, Small K, Marchini J (2016) Tensor decomposition for multiple-tissue gene expression experiments. Nature genetics 48: 1094–1100

Hothorn T, Lausen B (2003) On the exact distribution of maximally selected rank statistics. Computational Statistics & Data Analysis 43: 121–137

Huang Y, Sanguinetti G (2017) BRIE: transcriptome-wide splicing quantification in single cells. Genome biology 18: 123

Iorio F, Knijnenburg TA, Vis DJ, Bignell GR, Menden MP, Schubert M, Aben N, Goncalves E, Barthorpe S, Lightfoot H, Cokelaer T, Greninger P, van Dyk E, Chang H, de Silva H, Heyn H, Deng X, Egan RK, Liu Q, Mironenko T et al(2016) A Landscape of Pharmacogenomic Interactions in Cancer. Cell 166: 740–754

Jaakkola TS, Jordan MI (2000) Bayesian parameter estimation via variational methods. Statistics and Computing 10: 25–37

Khan SA, Virtanen S, Kallioniemi OP, Wennerberg K, Poso A, Kaski S (2014) Identification of structural features in chemicals associated with cancer drug response: a systematic data-driven analysis. Bioinformatics 30: i497–504

Kim M, Rai N, Zorraquino V, Tagkopoulos I (2016) Multi-omics integration accurately predicts cellular state in unexplored conditions for Escherichia coli. Nature communications 7: 13090

Klami A, Virtanen S, Leppaaho E, Kaski S (2015) Group Factor Analysis. IEEE transactions on neural networks and learning systems 26: 2136–2147

Lanckriet GR, De Bie T, Cristianini N, Jordan MI, Noble WS (2004) A statistical framework for genomic data fusion. Bioinformatics 20: 2626–2635

Leppäaho E, Kaski S (2017) GFA: exploratory analysis of multiple data sources with group factor analysis. Journal of Machine Learning Research 18: 1–5

Love MI, Huber W, Anders S (2014) Moderated estimation of fold change and dispersion for RNAseq data with DESeq2. Genome biology 15: 550

Lun AT, Bach K, Marioni JC (2016) Pooling across cells to normalize single-cell RNA sequencing data with many zero counts. Genome biology 17: 75

Macaulay IC, Haerty W, Kumar P, Li YI, Hu TX, Teng MJ, Goolam M, Saurat N, Coupland P, Shirley LM, Smith M, Van der Aa N, Banerjee R, Ellis PD, Quail MA, Swerdlow HP, Zernicka-Goetz M, Livesey FJ, Ponting CP, Voet T (2015) G&T-seq: parallel sequencing of single-cell genomes and transcriptomes. Nature methods 12: 519–522

Maloum K, Settegrana C, Chapiro E, Cazin B, Lepretre S, Delmer A, Leporrier M, Dreyfus B, Tournilhac O, Mahe B, Nguyen-Khac F, Lesty C, Davi F, Merle-Beral H (2009) IGHV gene mutational status and LPL/ADAM29 gene expression as clinical outcome predictors in CLL patients in remission following treatment with oral fludarabine plus cyclophosphamide. Annals of hematology 88: 1215–1221

Mazumder R, Hastie T, Tibshirani R (2010) Spectral Regularization Algorithms for Learning Large Incomplete Matrices. Journal of machine learning research : JMLR 11: 2287–2322

Meng C, Kuster B, Culhane AC, Gholami AM (2014) A multivariate approach to the integration of multi-omics datasets. BMC bioinformatics 15: 162

Meng C, Zeleznik OA, Thallinger GG, Kuster B, Gholami AM, Culhane AC (2016) Dimension reduction techniques for the integrative analysis of multi-omics data. Briefings in bioinformatics 17:628–641

Mertins P, Mani DR, Ruggles KV, Gillette MA, Clauser KR, Wang P, Wang X, Qiao JW, Cao S, Petralia F, Kawaler E, Mundt F, Krug K, Tu Z, Lei JT, Gatza ML, Wilkerson M, Perou CM, Yellapantula V, Huang KL et al(2016) Proteogenomics connects somatic mutations to signalling in breast cancer. Nature 534: 55–62

Mo Q, Wang S, Seshan VE, Olshen AB, Schultz N, Sander C, Powers RS, Ladanyi M, Shen R (2013) Pattern discovery and cancer gene identification in integrated cancer genomic data. Proceedings of the National Academy of Sciences of the United States of America 110: 4245–4250

Mohammed H, Hernando-Herraez I, Savino A, Scialdone A, Macaulay I, Mulas C, Chandra T, Voet T, Dean W, Nichols J (2017) Single-cell landscape of transcriptional heterogeneity and cell fate decisions during mouse early gastrulation. Cell reports 20: 1215–1228

Morabito F, Cutrona G, Mosca L, D’Anca M, Matis S, Gentile M, Vigna E, Colombo M, Recchia AG, Bossio S, De Stefano L, Maura F, Manzoni M, Ilariucci F, Consoli U, Vincelli I, Musolino C, Cortelezzi A, Molica S, Ferrarini M et al(2015) Surrogate molecular markers for IGHV mutational status in chronic lymphocytic leukemia for predicting time to first treatment. Leukemia research 39:840–845

Oakes CC, Seifert M, Assenov Y, Gu L, Przekopowitz M, Ruppert AS, Wang Q, Imbusch CD, Serva A, Koser SD, Brocks D, Lipka DB, Bogatyrova O, Weichenhan D, Brors B, Rassenti L, Kipps TJ, Mertens D, Zapatka M, Lichter P et al(2016) DNA methylation dynamics during B cell maturation underlie a continuum of disease phenotypes in chronic lymphocytic leukemia. Nature genetics 48: 253–264

Pierson E, Yau C (2015) ZIFA: Dimensionality reduction for zero-inflated single-cell gene expression analysis. Genome biology 16: 241

Plesingerova H, Librova Z, Plevova K, Libra A, Tichy B, Skuhrova Francova H, Vrbacky F, Smolej L, Mayer J, Bryja V, Doubek M, Pospisilova S (2017) COBLL1, LPL and ZAP70 expression defines prognostic subgroups of chronic lymphocytic leukemia patients with high accuracy and correlates with IGHV mutational status. Leukemia & lymphoma 58: 70–79

Queiros AC, Villamor N, Clot G, Martinez-Trillos A, Kulis M, Navarro A, Penas EM, Jayne S, Majid A, Richter J, Bergmann AK, Kolarova J, Royo C, Russinol N, Castellano G, Pinyol M, Bea S, Salaverria I, Lopez-Guerra M, Colomer D et al(2015) A B-cell epigenetic signature defines three biologic subgroups of chronic lymphocytic leukemia with clinical impact. Leukemia 29: 598–605

Remes S, Mononen T, Kaski S (2015) Classification of weak multi-view signals by sharing factors in a mixture of Bayesian group factor analyzers. arXiv preprint arXiv:151205610

Ritchie MD, Holzinger ER, Li R, Pendergrass SA, Kim D (2015) Methods of integrating data to uncover genotype-phenotype interactions. Nature reviews Genetics 16: 85–97

Seeger M, Bouchard G (2012) Fast variational Bayesian inference for non-conjugate matrix factorization models. In Artificial Intelligence and Statistics, pp 1012–1018.

Shen R, Olshen AB, Ladanyi M (2009) Integrative clustering of multiple genomic data types using a joint latent variable model with application to breast and lung cancer subtype analysis. Bioinformatics 25: 2906–2912

Singh A, Gautier B, Shannon CP, Vacher M, Rohart F, Tebutt SJ, Le Cao K-A (2016) DIABLO-an integrative, multi-omics, multivariate method for multi-group classification. bioRxiv: 067611

Soderholm S, Fu Y, Gaelings L, Belanov S, Yetukuri L, Berlinkov M, Cheltsov AV, Anders S, Aittokallio T, Nyman TA, Matikainen S, Kainov DE (2016) Multi-Omics Studies towards Novel Modulators of Influenza A Virus-Host Interaction. Viruses 8

Srivastava P (2002) Roles of heat-shock proteins in innate and adaptive immunity. Nature reviews Immunology 2: 185

Tenenhaus A, Philippe C, Guillemot V, Le Cao K-A, Grill J, Frouin V (2014) Variable selection for generalized canonical correlation analysis. Biostatistics 15: 569–583

Trachootham D, Alexandre J, Huang P (2009) Targeting cancer cells by ROS-mediated mechanisms: a radical therapeutic approach? Nature reviews Drug discovery 8: 579–591

Trojani A, Di Camillo B, Tedeschi A, Lodola M, Montesano S, Ricci F, Vismara E, Greco A, Veronese S, Orlacchio A (2012) Gene expression profiling identifies ARSD as a new marker of disease progression and the sphingolipid metabolism as a potential novel metabolism in chronic lymphocytic leukemia. Cancer Biomarkers 11: 15–28

Troyanskaya O, Cantor M, Sherlock G, Brown P, Hastie T, Tibshirani R, Botstein D, Altman RB (2001) Missing value estimation methods for DNA microarrays. Bioinformatics 17: 520–525

Vasconcelos Y, De Vos J, Vallat L, Reme T, Lalanne AI, Wanherdrick K, Michel A, Nguyen-Khac F, Oppezzo P, Magnac C, Maloum K, Ajchenbaum-Cymbalista F, Troussard X, Leporrier M, Klein B, Dighiero G, Davi F, French Cooperative Group on CLL (2005) Gene expression profiling of chronic lymphocytic leukemia can discriminate cases with stable disease and mutated Ig genes from those with progressive disease and unmutated Ig genes. Leukemia 19: 2002–2005

Virtanen S, Klami A, Khan S, Kaski S (2012) Bayesian group factor analysis. In Artificial Intelligence and Statistics, pp 1269–1277.

Wang B, Mezlini AM, Demir F, Fiume M, Tu Z, Brudno M, Haibe-Kains B, Goldenberg A (2014) Similarity network fusion for aggregating data types on a genomic scale. Nature methods 11: 333337

Westra H-J, Jansen RC, Fehrmann RS, te Meerman GJ, Van Heel D, Wijmenga C, Franke L (2011) MixupMapper: correcting sample mix-ups in genome-wide datasets increases power to detect small genetic effects. Bioinformatics 27: 2104–2111

Zenz T, Mertens D, Küppers R, Döhner H, Stilgenbauer S (2010) From pathogenesis to treatment of chronic lymphocytic leukaemia. Nature Reviews Cancer 10: 37–50

Zhao S, Gao C, Mukherjee S, Engelhardt BE (2016) Bayesian group factor analysis with structured sparsity. Journal of Machine Learning Research 17: 1–47

